# Mitochondrial metabolism and body condition of naturally infected sunfish (*Lepomis gibbosus*)

**DOI:** 10.1101/2022.09.19.508536

**Authors:** Vincent Mélançon, Sophie Breton, Stefano Bettinazzi, Marie Levet, Sandra A. Binning

## Abstract

Parasites can affect host behavior, cognition, locomotion, body condition and many other physiological traits. Changes to host aerobic metabolism are likely responsible for these parasite-induced performance alterations. Whole-organism metabolic rate is underpinned by cellular energy metabolism driven most prominently by the mitochondria. However, few studies have explored how mitochondrial enzymatic activity relates to body condition and parasite infection despite being a putative site for metabolic disruptions related to health status. We studied correlations among natural parasite infection, host body condition and the activity of key mitochondrial enzymes in target organs from wild-caught pumpkinseed sunfish (*Lepomis gibbosus*) to better understand the cellular responses of fish hosts to endoparasite infection. Enzymatic activities in the gills, spleen, and brain of infected fish were not significantly related to parasite infection or host body condition. However, the activity of cytochrome C oxidase, an enzyme involved in oxidative phosphorylation, in fish hearts was higher in individuals with lower body condition. Activities of citrate synthase, complexes I and III and carnitine palmitoyltransferase were also significantly different among organ types. These results provide preliminary information regarding the likely mitochondrial pathways affecting host body condition, the maintenance energetic requirements of different organs and their specific dependency on particular mitochondrial pathways. These results help pave the way for future studies on the effects of parasite infection on mitochondrial metabolism.

## Introduction

Most wild animals are infected with parasites (Dobson et al. 2008). Parasites are organisms living in or on another specie, the host, that they exploit for resources (shelter and/or energy) (Lewin 1982). However, quantifying parasite-induced damages and/or costs to hosts is often difficult (Araújo et al. 2003; Leung and Poulin 2008). In addition, parasites are often much smaller than their host, making infection difficult to detect and, thus, easy for researchers to overlook (Marcogliese 2004). This oversight can be problematic considering that many parasites have important effects on host physiology and behavior (Lagrue and Poulin 2015; Timi and Poulin 2020). For instance, body condition, the relationship between an animal’s mass and body size, is used in many taxa as a proxy of individual health status (Jakob et al. 1996). However, when infected individuals are weighed, the mass of their parasites is inevitably included in this measure. This can lead to an overestimate of an individual’s body condition in heavily infected individuals if host mass is not corrected for (Lagrue and Poulin 2015; Timi and Poulin 2020). Thus, considering parasite infection in eco-physiological studies is critical to ensure accurate estimates of many traits of interest.

Host performance capacity (i.e., the ability of an organism to execute fitness-enhancing behaviors) has recently been identified as a target of direct or indirect manipulation by parasites (Poulin 2010; McElroy and De Buron 2014; Binning et al. 2017; Timi and Poulin 2020). For example, bridled monocle bream (*Scolopsis bilineatus*) infected by the ectoparasitic isopod (*Anilocra nemipteri*) experience decreased aerobic capacity and maximum swimming speed compared with uninfected fish (Binning et al. 2013). Similarly, pumpkinseed sunfish (*Lepomis gibbosus*) heavily infected with bass tapeworms (*Proteocephallus ambloplitis*) display reduced responsiveness and standard and maximum metabolic rates suggesting that parasites may have an effect on host aerobic physiology and escape performance (Guitard et al. 2022). Parasite infection can also increase host energy demand by triggering the costly activation of the host immune system or by causing tissue damage or other parasite-induced pathologies (Ots et al. 2001; Freitak et al. 2003). For example, house sparrows (*Passer domesticus*) injected with lipopolysaccharide (LPS) from *Escherichia coli*, an endotoxin that induces an immune response in most vertebrates, show decreased activity, feeding rates and reproductive success (Bonneaud et al. 2003). There is also nestling Mediterranean blue tits (*Parus caeruleus L*.) that, when infected with *Protocalliphora* ectoparasites, experience decreased aerobic competence (Simon et al. 2004) or white cabbage butterflies (*Pieris brassicae L*.) that, when implanted with nylon to mimic a parasitic infection, have increased metabolic rates (Freitak et al. 2003). Despite the growing body of work linking infection to altered energetics, the mechanisms underlying altered host metabolic activities during infection are not well understood.

In addition to energetics, host body condition tends to be negatively related to parasite intensity (Lemly and Esch 1984; Sánchez et al. 2018). Body condition reflects the amount of stored lipids in an organism. Thus, individuals with a lower condition index likely have lower energy reserves than conspecifics with a higher condition index. For instance, intracellular parasites like *Toxoplasma gondii* can exploit the host lipophagy machinery to access the fatty acids necessary to boost their own proliferation, at the expenses of host lipid reserve and body condition (Charron and Sibley 2002; Caffaro and Boothroyd 2011; Pernas et al. 2018). Similarly, in female springbok (*Antidorcas marsupialis*) and horses (*Equus ferus*), body condition is negatively correlated with increasing parasite loads (Turner et al. 2012; Debeffe et al. 2016). A negative correlation between infection and body condition also occurs in juvenile bluegill sunfish (*Lepomis macrochirus*) infected by trematodes (*Uvulifer ambloplitis*) (Lemly and Esch 1984) and in icefish (*Chionodraco hamatus*) infected with flatworms (helminths) (Santoro et al. 2013). The altered body condition and metabolic rate observed in infected hosts may be linked, but once again, the underlying mechanisms are not well explored in the literature.

The study of these underlying mechanisms starts with the mitochondria, which are key regulators of cellular energy metabolism. These organelles produce most of the cells’ energy (ATP) through a process known as oxidative phosphorylation (OXPHOS). The morphology of mitochondria is known to be affected by intracellular parasites; a process essential to promote parasite growth. For example, mitochondria elongate following infection by *Toxoplasma gondii* (Pernas et al. 2018). In contrast, they fragment following infection by *Listeria monocytogenes* (Stavru et al. 2011) or *Vibrio cholerae* (Suzuki et al. 2014a). However, how mitochondria respond to extracellular parasites is less well understood. OXPHOS is one of the primary users of oxygen (Capaldi 1990; Nath and Villadsen 2015) and some studies suggest a lowered OXPHOS activity in response to parasite infection (Mills et al. 2017; Hunter-Manseau et al. 2019). For example, when mounting an immune response to infection, the metabolism sometimes shifts toward the anaerobic glycolytic pathway (Escoll et al. 2019; Hortová□ Kohoutková et al. 2021). Mitochondria can also change their fuel of choice in response to different stressors (Stanley et al. 2014). For example, mice infected by *Toxoplasma gondii* shift towards lipid metabolism in response to infection to prevent rapid parasite growth (Pernas et al. 2018). This “metabolic defense” involves mitochondria dynamics, specifically fusion and grouping around the *Toxoplasma* vacuole, together with an enhanced fatty acid oxidation, in order to compete with the parasite for key resources (Pernas et al. 2018) The metabolism of lipids, precisely fatty acid oxidation may, thus, increase with parasite infection. This could also explain why host body condition tends to be lower when individuals have a higher parasite intensity since a higher lipid metabolism activity decreases the amount of stored lipids (Pernas et al. 2018).

Physiological studies should also consider different organ systems because they may have varying responses to stressors, including parasite infection. For example, the brain is known for its high energetic demand (Magistretti and Allaman 2013) and uses most of the glucose, amino acids and monocarboxylates available to an organism. The brain uses primarily the OXPHOS and the tricarboxylic acid (TCA) cycle mitochondrial metabolic pathway, which can vary in response to environmental stressors (Soengas and Aldegunde 2002). Some evidence suggests that infection with the trematode parasite (*Euhaplorchis californiensis*) in the brains of the California killifish (*Fundulus parvipinnis*) affects brain enzyme activities (Nadler et al. 2021). In comparison, the spleen is a small organ that is primarily used in immune responses, iron recycling (hematopoiesis) and blood filtration (Fänge and Nilsson 1985). The normal energetic demand of this organ is not high, but can greatly increase when an immune response is initiated (Bronte and Pittet 2013). In fishes, the heart and the gills are both important for the circulation and the delivery of oxygen throughout the body (Laurent et al. 1983). The energetic demand of the heart and the gills may also vary in response to stress. For example, gilthead seabream (*Sparus auratus*) exposed to cortisol, a stress hormone, show different gill enzymatic activities and mitochondrial pathway prioritization (Laiz-Carrión et al. 2002). Parasite infection can also induce stress in hosts (O’Dwyer et al. 2020) suggesting that parasites may indirectly affect enzymatic activity in gills and other tissues. These relationships have not been well explored to date.

The objective of this study is to measure the mitochondrial metabolism of a freshwater fish naturally parasitized with helminth endoparasites and to explore relationships among infection intensity, enzymatic activity, and body condition in four key organs. We hypothesize that the activity of key enzymes involved in the OXPHOS, the fatty acid metabolism and the TCA cycle are related to parasite intensity in naturally infected individuals, specifically that their activity increases with higher parasite intensities.

## Methods

### Study system

Pumpkinseed sunfish (*Lepomis gibbosus*) are small, common, freshwater fish native to Eastern North America. In lakes around the Laurentian region in Quebec, they are often the second intermediate host of several parasitic helminths including the cestode *Proteocephalus ambloplitis* (bass tapeworm), and the trematode *Clinostomum marginatum* (yellow grub). Pumpkinseed sunfish are also second intermediate hosts of the trematodes *Uvulifer sp*., and *Apophallus sp*., that form melanized cysts, which are visible on the fish’s body surface and fins and can be easily quantified by researchers. This visible sign of parasitism, referred to colloquially as black spot disease, allowed us to select fish across a gradient of infection upon capture. These parasites have complex lifecycles involving gastropods or copepods, fish and piscivorous birds or fishes as hosts. Trematode infection in fish occurs when the parasite cercaria are released from a gastropod and penetrate the muscles and fins of fish. The cercaria develop into metacercaria forming a cyst that is melanized by the host’s immune system leading to a visible black spot (Fischer and Freeman 1969; Berra and Au 1978). Once encysted, trematode metacercaria are metabolically inactive. However, heavy infections have been associated with altered behavior, reduced body condition and lipid content, and increased overwinter mortality in sunfish, suggesting a high energetic cost of infection (Lemly and Esch 1984; Timi and Poulin 2020). Cestode infection occurs when fish consume infected copepods. The cestode final hosts are other fishes like the small and largemouth bass (*Micropterus dolomieu* and *M. salmoides*) that become infected after ingesting infected sunfish. In sunfish, cestodes are found in the digestive tract and abdominal cavity, more precisely on the liver and the spleen. There, they cause important lesions and possible organ failure. Thus, they are suspected to have a large effect on the metabolism of the fishes by affecting the amount of nutrients received by the host and, perhaps, indirectly affecting mitochondria (Hugghins 1959; Margolis and Arthur 1979).

### Fish sampling

Pumpkinseed sunfish (*Lepomis gibbosus*) were collected using baited minnow traps from Lake Cromwell near the Station de Biologie des Laurentides (SBL) in Quebec, Canada (DD: 45.98861, −74,00585). Traps were set for 30 to 60 minutes before being pulled in and screened for fish. In total, 22 fish measuring between 90mm to 120mm in total length were collected across a gradient of visible black spot infection, which was assessed by the researcher (VM) upon capture and later confirmed by the quantification of black spots on dead specimen in the lab. Selected fish were immediately euthanized in an overdose of 10% eugenol (clove-oil) solution (4 mL/L of water) and packed on ice in coolers. Fish that were not selected because they fell outside of the targeted size range or infection level were immediately released. Fish were quickly brought to the SBL and frozen at −20°C before being transported to UdeM’s Complexe des sciences laboratory facilities in refrigerated coolers for dissection and enzymatic assays.

### Fish dissection and body condition calculation

Each specimen was thawed and dissected on ice to maintain the enzymatic capacities of the tissues. First, fish total length was measured using a 0-150mm digital caliper. The total wet-weight was measured using an electronical balance (MSE225S, Sartorius Weighing Company). Second, the heart, spleen and brain were extracted and kept on ice in 5mL cryogenic tubes. Gill filaments were dissected out on ice using a dissecting microscope (Stemi DV4, Zeiss) and also kept on ice in cryogenic tubes once dissected. Third, fish were screened for parasites. Cestodes were extracted out of the fish and counted. Trematodes causing black spot disease were counted on the left side of fish only to avoid double counting parasites on unpaired fins, then doubled. Previous work suggests that there is no significant difference in the number of parasites found on the left vs the right side of the fish (Binning, Unpublished data). Lastly, the corrected weight of the fish (fish mass excluding cestode parasites) was measured by weighing the fish and tissues once all cestodes were removed. The weight of individual blackspot cysts is negligeable and thus not corrected for. The corrected fish mass was used to calculate the density of both types of parasites (trematodes and cestodes) for each fish by dividing the number of parasites in each fish by the corrected mass (Lagrue and Poulin 2015).

Le Cren’s body condition index was calculated for each fish. This index uses the log relation between the corrected mass and the total length of the fish to estimate a slope and an intercept for the population. These values are then used to measure the body condition of each individual that is centered around the population mean value of 1 (Cren 1951; Wuenschel et al. 2019).

### Enzymatic assays

Mitochondrial enzymatic activities were measured along a natural gradient of infection with the goal to explore possible correlations between the mitochondrial metabolism and parasite infection. These analyses were performed on the brain, heart, gills and spleen because of their relevance in various physiological systems. Few enzymes were selected to assess multiple mitochondrial pathways, more precisely the oxidative phosphorylation, the tricarboxylic acid cycle, and the lipid metabolism because of the possible effects of parasitism on them.

Once the organs were extracted from the fish, they were diluted 20 times their wet-weight in a 5 mL cryogenic tube with a 100 mM potassium phosphate buffer, 20 mM ethylene diamine tetraacetic acid (EDTA), pH 8,0 (modified protocol from Hunter-Manseau et al. 2019). Then, they were homogenized using a polytron (PT1200 E) three times for 5 seconds with a rest time on ice of 30 seconds between each step. The homogenized samples were stored at −80°C.

Enzymatic assays were performed at room temperature in 96-well plates using a microplate reader (Mithras LB 940, Berthold Technologies). Cytochrome C oxidase (CCO) and electron transport system (ETS – complexes I and III) were chosen because of their importance in the OXPHOS (Mills et al. 2017; Ryan et al. 2021). To explore the relationships between parasitism and lipid metabolism, carnitine palmitoyltransferase (CPT) was selected due to its role in transferring a carnitine group to fatty acids thus allowing their entry in the mitochondrial matrix (Schulz 1991; Hiltunen et al. 2010; Kastaniotis et al. 2017). An enzyme from the TCA cycle, citrate synthase (CS), was also chosen as a quantitative enzyme marker of mitochondria content (Mills et al. 2017; Ryan et al. 2021).

All assays were run in duplicate. Enzyme activities were estimated using the Beer-Lambert law (Thibault et al. 1997), corrected for the dilution factor and normalized for the amount of proteins of each sample. All the reagents came from Sigma-Aldrich chemicals (St-Louis, USA).

### Enzymatic activity

#### Cytochrome c oxidase

(CCO, EC 7.1.1.9): CCO activity was measured at 550 nm over 5 minutes following the oxidation of cytochrome c (ε=18.5 mM^−1^·cm^−1^) using a 100 mM potassium phosphate buffer containing 0.05 mM cytochrome C from equine heart, 4.5 mM sodium dithionate (DTT) and 0.03 % triton X 100, pH 8.0. Control reactions were performed by adding 10 mM of sodium azide to the sample and background activities were done by omitting DTT then deducted to the activity of the assay. The reaction solution was bubbled for 5 minutes and the absorbance ratio of 550nm/565nm was determined. If the ratio was higher than 9, the solution was used (Thibault et al. 1997; Hunter-Manseau et al. 2019).

#### Electron transport system

(ETS, EC 7.1.1.2 and 7.1.1.8): ETS activity (mitochondrial complexes I + III) was assessed at 490 nm over 6 minutes by measuring the reduction of p-iodonitrotetrazolium violet (INT, ε =15.9 mM^−1^·cm^−1^) using a 100 Mm potassium phosphate buffer with 2 mM INT and 0.85 mM NADH, pH 8.5. Background and specificity of the reaction were calculated by running a control without tissue and a control without NADH (Hunter-Manseau et al. 2019).

#### Carnitine palmitoyltransferase (CTP, EC 2.3.1.21)

CPT activity was assessed at 412 nm over 5 minutes by measuring the conversion of 5,5’ dithiobis-2-nitrobenzoic acid (DTNB) into TNB (ε=14.15 mM^−1^·cm^−^1) using a 75 mM tris-HCl buffer, 1.5 mM EDTA with 0.25 mM DTNB, 0.035 mM PalmitoylCoA and 2 mM L-carnitine, pH 8.0. Control reactions were performed by omitting the sample (Thibault et al. 1997).

#### Citrate synthase (CS, EC 2.3.3.1)

CS activity was measured at 412nm following the conversion of 5,5’ dithiobis-2-nitrobenzoic acid (DTNB) into TNB (ε=14.15 mM^−1^·cm^−^1) over 6 minutes. The reaction solution consisted of 100mM imidazole-HCl buffer with 0.1 mM DTNB, 0.1 mM acetyl CoA and 0.15 mM oxaloacetate, pH 8.0. A background without oxaloacetate was first performed followed by the final assay (Hunter-Manseau et al. 2019).

Enzyme activities were normalized by total protein content (mg mL^−1^) measured through the bicinchoninic acid method (Smith et al. 1985) and are expressed as U mg protein^−1^ where U represents 1 μmol of substrate transformed to product in 1 min.

### Statistical analyses and measures

#### Parasites and host body condition

– To explore the relationship between parasite infection and host body condition, generalized linear models (GLM) with a Gamma error distribution were used to meet model assumptions. Two models were tested: 1) host body condition as a function of cestode density and 2) host body condition as a function of trematode density. Both parasites have different ecologies and likely have different effects on their hosts. Their intensities are also not strongly correlated in our sample (R^2^=0,35). However, we considered them separately in our analyses.

#### Parasites and enzymatic activity

– To explore the relationship between parasite infection and enzymatic activity, linear mixed models (LMM) were used. Response variables were boxcox transformed to meet model assumptions. For each of the four enzymes tested (CCO, CS, ETS, CPT), two models were run: 1) enzymatic activity as a function of cestode density and 2) enzymatic activity as a function of trematode density. Organ type (heart, spleen, gills, brain) was included as a fixed factor and body condition as a covariable in both models. Full models included a three-way interaction between parasite density, organ type and body condition with fish ID as a random effect. Simplified models were selected based on their AICc: the model with the lowest AICc was kept. At first, both model 1) and 2) were tested within the same model without interaction between these two three-way interaction models for each enzyme. The AICc showed that this model was not the most appropriate for all enzymes and was therefore discarded. For CS, CCO and CPT, the selected model included only body condition, organ type and their interaction as factors. For ETS, the model with cestode density, organ type and their interaction was chosen. When interactions between factors and covariables were not significant, interactions were removed from the model. All statistics were performed using the lmer, glm and ggplot functions in R 4.1.0 (R Development Core Team 2022).

## Results

### Parasites and host body condition

Fish measured in this study had a mean total length of 103 mm (91.9 – 121.5 ± 8.9 mm; min-max ± SE). Mean fish wet-weight was 21.5 g (15.4 – 36.4 ± 5.3 g) and the mean parasite mass-adjusted wet-weight was 20.6 g (14.7 – 35.6 ± 5.4 g). Fish body condition index ranged from 0.82 to 1.19. Trematode density ranged between 0.06 and 29.4 (10.4 ± 14.2; mean ± SE) trematodes g^−1^. Individuals carried as few as 2 trematodes and as many as 518 trematodes (median = 10.1 trematodes). For cestodes, parasite density was lower with a minimum of 0.3 and a maximum of 11.1 (median = 2.8 ± 1.5) cestodes g^−1^. The cestode intensity ranged from 4 to 210 cestodes per fish (median = 2.1 cestodes). Even though there appeared to be a negative correlation between fish parasite density and host body condition for both types of infection (figure S1), this effect was not statistically significant (cestodes R^2^=0.03; trematodes R^2^=0.11; both p-value > 0.1).

### Parasites and host enzymatic activity

For CCO enzyme activity, neither model containing parasite density was significant (see table S1) (figure 1A and 2A). In the model containing host body condition and organ type, there was a significant interaction between the organ type and fish body condition (F-value = 3.00; p-value = 0.038). Increased body condition was correlated with a decrease in CCO activity for the heart (figure 3A) (t-value = −2.27; p-value = 0.027), but not other organs.

**Figure 1:**
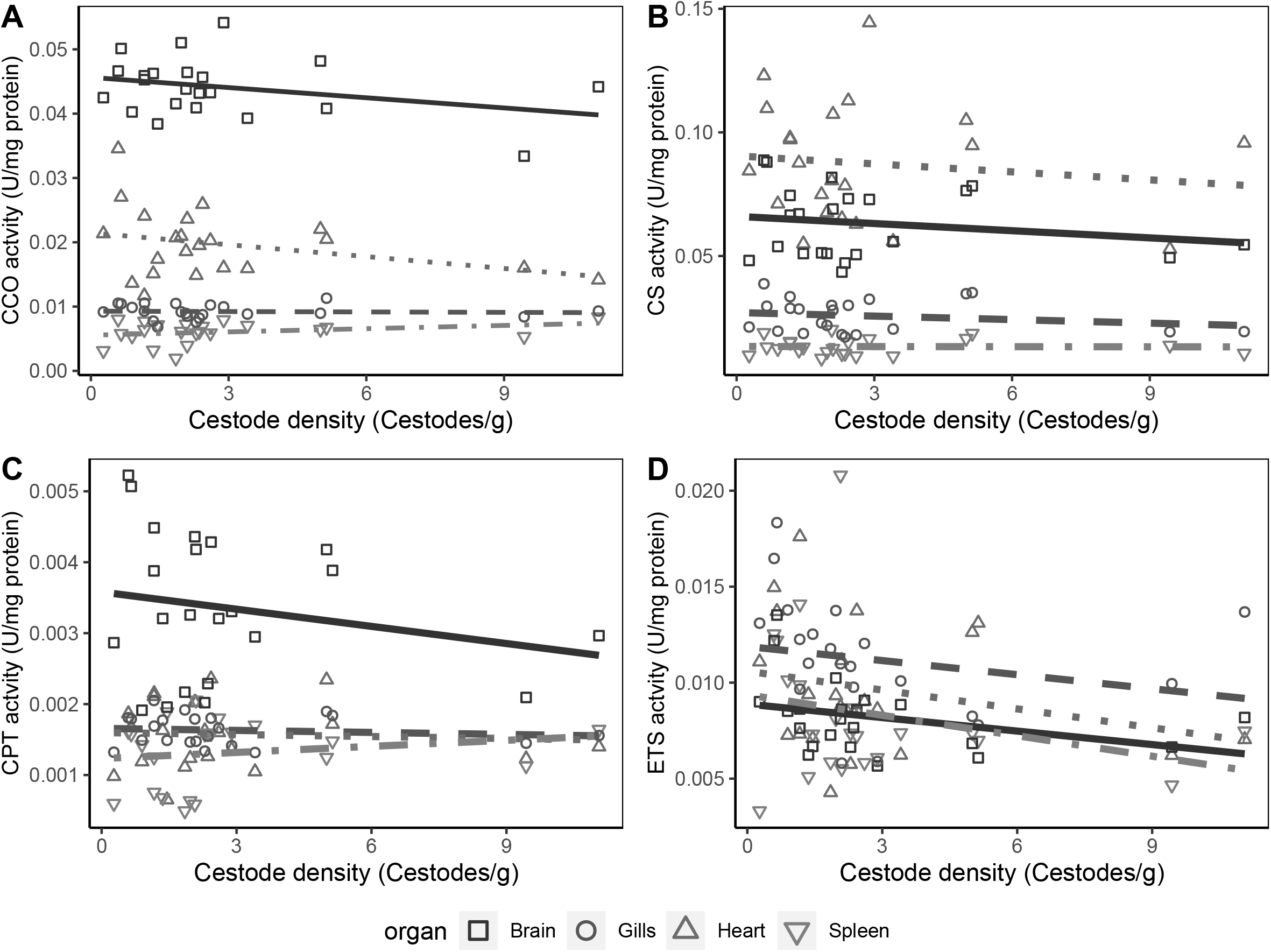
Relationships among the cestodes parasite infection and the enzymatic activity of all selected enzymes. A) The enzymatic activity (U mg^−1^ protein) as a function of the cestode density of cytochrome c oxidase (CCO). B) The enzymatic activity (U mg^−1^ protein) as a function of the cestode density of citrate synthase (CS). C) The enzymatic activity (U mg^−1^ protein) as a function of the cestode density of carnitine palmitoyltransferase (CPT). D) The enzymatic activity (U mg^−1^ protein) as a function of the cestode density of the electron transport system (ETS – mitochondrial complexes I and III).

**Figure 2:**
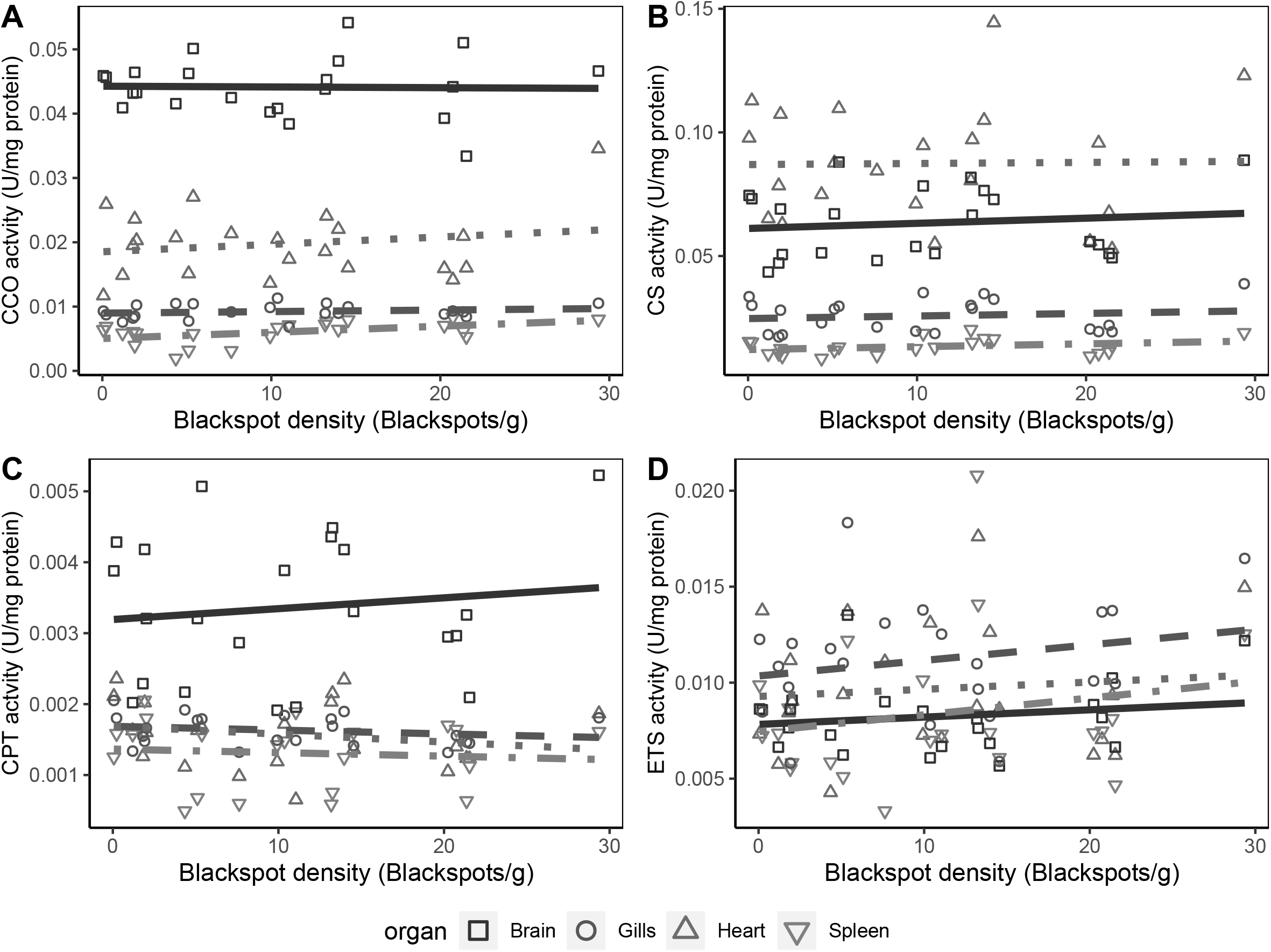
Relationship between the black spot parasite infection and the enzymatic activity of all enzymes. A) The enzymatic activity (U mg^−1^ protein) as a function of the black spot density of cytochrome c oxidase (CCO). B) The enzymatic activity (U mg^−1^ protein) as a function of the black spot density of citrate synthase (CS). C) The enzymatic activity (U mg^−1^ protein) as a function of the black spot density of carnitine palmitoyltransferase (CPT). D) The enzymatic activity (U mg^−1^ protein) as a function of the black spot density of the electron transport system (ETS – mitochondrial complexes I and III).

**Figure 3:**
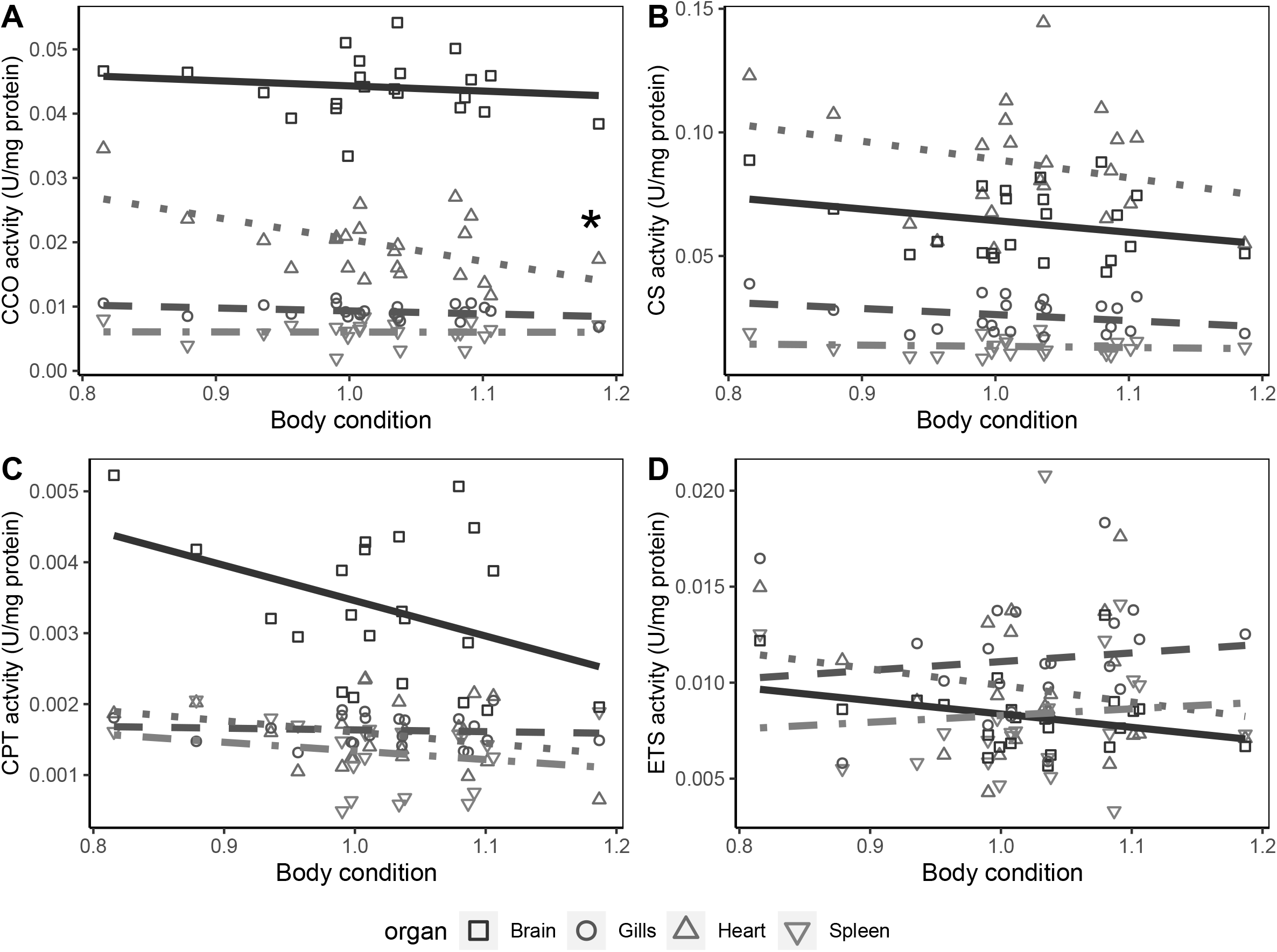
Relationship between the body condition and the enzymatic activity of all selected enzymes. A) The enzymatic activity (U mg^−1^ protein) as a function of the body condition of cytochrome c oxidase (CCO). B) The enzymatic activity (U mg^−1^ protein) as a function of the body condition of citrate synthase (CS). C) The enzymatic activity (U mg^−1^ protein) as a function of the body condition of carnitine palmitoyltransferase (CPT). D) The enzymatic activity (U mg^−1^ protein) as a function of the body condition of the electron transport system (ETS – mitochondrial complexes I and III). (* = p < 0.05).

A different situation was seen with CS activity. There was no significance in the models that included the parasite density and organ type (see table S1) and no significant interaction between the body condition and the organ (p-value = 0.42) in the model including both factors (figure 1B–2B–3B). However, a significant difference between organs was detected (p-value = < 2e^−16^) in the model including body condition and organ type without interactions. These differences are between the spleen (0.013 ± 0.00071 U mg protein^−1^) and the brain (0.063 ± 0.003 U mg protein^−1^) (p-value < 2e^−16^); between the heart (0.087 ± 0.0051 U mg protein^−1^) and the gills (0.026 ± 0.0014 U mg protein^−1^) (p-value < 2e^−16^); between the spleen and the heart (p-value = < 2e^−16^); between the brain and the gills (p-value < 2e^−16^); between the heart and the brain (p-value< 3.52e^−12^); and between the gills and the spleen (p-value< 1,04e^−12^) (figure 4A).

**Figure 4:**
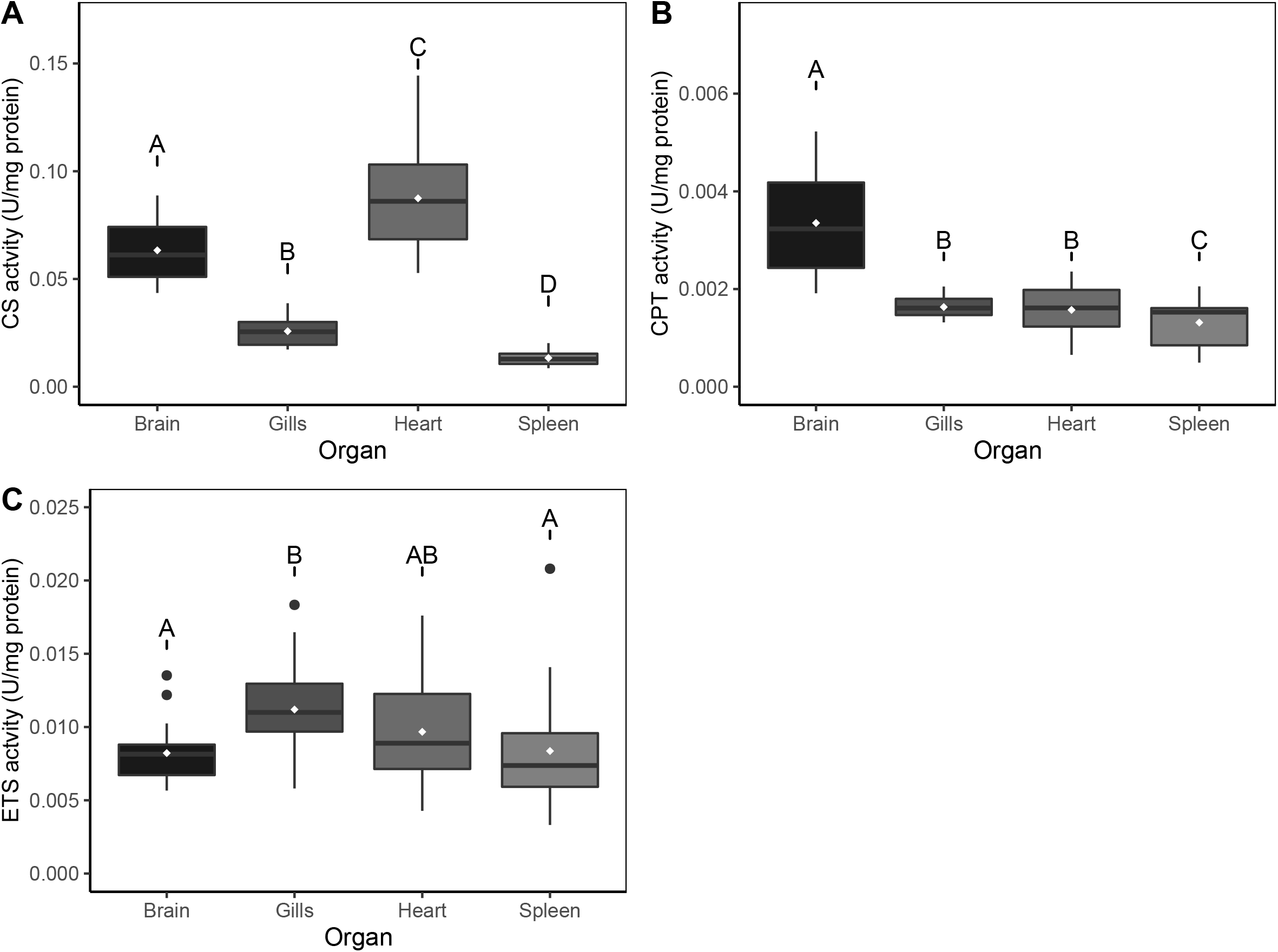
Differences of enzymatic activity between organs. A) The boxplot representation of the differences of CS activity (U mg^−1^ protein) between all selected organs. B) The boxplot representation of the differences of CPT activity (U mg^−1^ protein) between all selected organs. C) The boxplot representation of the differences of ETS activity (U mg^−1^ protein) between all selected organs. The with diamonds represent the mean enzymatic activity of each organ.

None of the previously mentioned models including parasite density were found to be significant in the case of CPT (see table S1), but trends were present for body condition (p-value = 0.062) (figure 1C–2C–3C). Like in the case of CS, the organ type had a significant effect (p-value = < 2e^−16^) with the brain having a higher enzymatic activity (0,0034 ± 0,00021 U mg protein^−1^) compared to the gills (0,0016 ± 0,000046 U mg protein^−1^) (p-value = 4,41e^−14^), the heart (0,0016 ± 0,000010 U mg protein^−1^) (p-value = 3,83e^−15^) and the spleen (0,0013 ± 0,00010 U mg protein^−1^) (p-value = < 2e^−16^). Also, the spleen had a significantly different CPT activity compared to the gills (p-value = 0,0087) and to the heart (p-value = 0,041) (Figure 4B). These results were obtained using the statistical model including only body condition and organ type without their interaction.

For ETS, there was no significant relationship between any of the tested factors and enzymatic activity for models with either cestode density or black spot density (see table S1) (figure 1D–2D–3D) except for the organ type (p-value= 0,00078) in the model including cestode density and organ type without interactions. Cestode density approached significance (p-value = 0.093) and the gills ETS activity (0,011 ± 0,00065 U mg protein^−1^) was found to be significantly different of the brain activity (0,0082 ± 0,00041 U mg protein^−1^) (p-value = 0,00051) and the spleen (0,0084 ± 0,00082 U mg protein^−1^) (p-value= 0,00031) (Figure 4C).

## Discussion

The goal of this study was to explore the mitochondrial metabolism of parasitized *L. gibbosus*. We found no significant relationship between parasite density and host mitochondrial enzymatic activity in any of the tested enzymes in the four organs we studied. However, there was a significant negative relationship between body condition and CCO activity of the heart. Indeed, the heart’s activity was higher when an individual had lower body condition. The activity of CS varied significantly among organs with brain and heart activity being significantly higher than the spleen, and heart activity also being higher than that of the gills. The activities of ETS and CPT were also found to be significantly different between organs: the gills and the brain had the highest activity respectively.

### Parasites and host body condition

We found no correlation between parasite density and host body condition, despite a negative trend with both types of parasites (figure S1). Although parasites tend to be negatively related to host body condition, some studies have also found positive or no relationships between these variables (Sánchez et al. 2018). Our results slightly contrast with those reported by Lemly and Esch (1984), who found a significant negative relationship between condition and infection with similar trematode parasites (*Uvulifer ambloplitis*) in bluegill sunfish (*Lepomis macrochirus*). This discrepancy may be because Lemly and Esch (1984) explored these relationships in a semi-controlled experimental set-up (cages placed in a lake), as opposed to our wild-caught fish. In our case, it is impossible to know when parasite infections occurred in hosts, and thus whether fish were experiencing acute stages of infection that are more likely to be associated with increased energetic costs (Lemly and Esch 1984). Given that the cestodes we study are known to damage the liver, an important glycogen reserve, we expected to find a significant negative relationship between parasite infection and host body condition. Heavy damage to the liver may result in loss of function and host fish resorting to other stored metabolic substrates (Santoro et al. 2013). For example, Slavik et al. (2017) found that *Cyprinus carpio* infected by glochidia of the Chinese pond mussel (*Sinanodonta (Anodonta) woodiana*) had an altered energy metabolism that could not be explained by variation of movement activities but by biochemical alterations of the liver and other organs. Although it would have been interesting to measure liver enzymatic activities directly, this is difficult since the liver tissue was often so heavily damaged in our population of fish that uninfected tissue was difficult to discriminate and collect properly with our dissection technique. Protocols to collect intact liver tissue from this study system are currently being developed and offer an exciting avenue for future research (Mélançon, Unpublished data). Also, while we could visually assess infection with black spot disease prior to collection, infection with cestodes could only be determined upon dissection. Since parasites tend to aggregate in hosts (Shaw and Dobson 1995), our collection protocol made it difficult to sample many fish heavily infected with cestodes.

### Parasites and host mitochondrial enzymatic activity

Although no statistically significant relationship was detected, cestode density tended to be negatively related to mitochondrial enzymatic activity whereas trematode density tended to be positively related to enzymatic activity (figures 1 and 2). These trends are interesting, and merit further investigation. For instance, cestodes are metabolically active while in the host, and may have a bigger impact on aerobic capacity, swimming speed and other fitness related traits. Indeed, Guitard et al. (2022) found negative correlation between host fish standard metabolic rate and cestode infection suggesting reduced metabolic activity during heavy infection. Trematodes, on the contrary, are dormant once encysted, but activate the host’s immune system during the encystment process. This process is related to increased metabolic rates in hosts (Lemly and Esch 1984), which suggests increased mitochondrial activity as well. Experimental infection with trematodes could help capture the acute phase of this infection when metabolic activities of hosts are likely the most affected.

### Enzymatic activity and host body condition

Our results showed that CCO activity is correlated with a decreasing body condition index but only in one organ, the heart (figure 3A). CCO is the mitochondrial enzyme involved in the transfer of electrons to oxygen, the final electron carrier in aerobic cellular respiration. CCO is believed to be the front-runner for mitochondrial oxidative metabolism (Capaldi 1990; Lemieux and Blier 2022). Higher enzymatic activity in the heart means increased overall capacity for energy production and muscular activity. Since the heart is responsible for the circulation of the blood throughout the body, increased energy production may lead to increased blood circulation to tissues. It is possible that fish in poor body condition experience increased nutrient demand, and thus have greater heart activity. Alternatively, heart morphology may be modified by infection. In cold environments or in food deprived state, fish hearts can undergoe hypertrophy or hyperplasia in order to counter the decreased contractility caused by the environment temperature that affects the stroke volume of hearts (Gamperl and Farrell 2004). A similar response may occur following the biotic stress of parasite infection (Holmstrup et al. 2010). This could explain why an individual with a lower body condition index has a higher heart CCO activity. Including heart somatic index (i.e., heart mass expressed as a percentage of body wet mass) or another metric of heart morphology would be useful in the future to quantify this phenomenon and confirm the hypertrophy or hyperplasia hypothesis in response to a parasite infection (Driedzic et al. 1996). In general, we tend to see a non-significant negative trend between enzymatic activity and body condition for all organs. Some of these trends are close to being significant especially in CPT’s case, justifying increasing the sample size to clarify these trends.

### Enzymatic activity and organ type

Although activities of most tested enzymes did not change in response to parasite or body condition, they all differed significantly between organs. One notable example is citrate synthase, an enzyme of the tricarboxylic acid cycle widely used as a proxy of mitochondria density. The heart and the brain had a significantly higher CS activity than other organs possibly indicating a higher mitochondria density (figure 4A). This coincides with the fact that the heart and the brain are two highly energy depending organs. Increased TCA activity and/or mitochondrial density leads to more reducing equivalent feeding the OXPHOS pathway, which may further increase ATP production. The heart has high baseline energetic demand (Laurent et al. 1983). By undergoing hypertrophy or hyperplasia in response to physiological stress induced by parasite infections, it is likely that hearts require an increase in CS activity and/or increased number of mitochondria. These relationships should be further explored. Another interesting observation was carnitine palmitoyltransferase activity, which was highest in the brain (Figure 4B). This organ has stored lipid that can be used by neuronal cells to produce ATP (Soengas and Aldegunde 2002). By having a higher CPT activity, it shows that the brain potentially tends to consume more of its lipid reserve then other organs, possibly to compensate for a lack of substrates coming from the digestive system, a consequence of parasitism.

The spleen is a small organ with a low energetic demand when not mounting an immune reaction, and we found low activity in this organ for all tested enzymes. More research is needed to compare the enzymatic activity of spleen from fish with active immune systems. A study comparing fish injected with an immune stimulant such as LPS and/or fish that are infected experimentally could help correlate immune response to parasite infections.

### Conclusion

When studying natural populations, multiple biotic and abiotic factors should be considered (Holmstrup et al. 2010). Parasite infections are important biotic stressors that are overlooked compared with abiotic factors such as temperature, pH, and hypoxia despite their potential physiological costs. Thus, overlooking parasitic infection in physiology studies can be problematic (Suzuki et al. 2014b; Timi and Poulin 2020). Our study represents the first attempt to link parasite infection with changes in mitochondrial bioenergetics in four different organs of freshwater fish characterized by a gradient of parasite infection. Our results suggest that cytochrome c oxidase activity in the heart might be linked to body condition and that citrate synthase, electron transport system and carnitine palmitoyltransferase activities differs significantly between organs. Even though there were no significant relationships between fish parasite load and the mitochondrial metabolism, interesting trends help pave the way towards a clearer understanding of cellular mechanisms underlying metabolic alterations experienced during infection, something not currently well explored in the literature. Additional sampling to obtain a broader range of cestode infection would help reveal trends across ecologically relevant infection gradients. Further studies should continue to pursue these questions across a greater range of organs (i.e. liver, muscle, gonads) and enzymes (e.g. enzymes of the glycolytic pathway such as lactate dehydrogenase and anti-oxidant enzymes such as catalase and glutathione peroxidase, etc.) to deepen our mechanistic understanding of the role of parasites on host physiology in natural systems.

## Acknowledgments

We acknowledge the traditional lands of the Kanien’kehà:ka, Omàmiwinini and Anishinabewaki First Nations on which the field and laboratory work for this project took place. We thank Gabriel Lanthier for logistic support at the SBL and Amélie Papillon for help with fish collection. We also thank Shaun Killen, Ariane Côté, Maryane Gradito, Jérémy De Bonville and Emmanuelle Chrétien for helpful discussions and/or comments on earlier versions of this manuscript. Funding was provided by a Seed grant for Innovative Research Projects from the Groupe de Recherche Interuniversitaire en Limnologie (SAB, SB, SK). SAB and SB are also supported by the Canada Research Chair program. Animals were collected following protocols approved by UdeM’s Comité de déontologie de l’expérimentation sur les animaux (protocol number 21-028) and collection permits were granted by the Ministère des Forêts, de la Faune et des Parcs (Permit number: SEG#2021-05-18-1833-15SP).

## Electronic supplementary material

*Figure S1: Relationship between parasite density and host body condition*. Le Cren’s body condition in relation to the density of cestodes (A) or trematodes (B). Grey zones represent the 95% confidence interval.

*Table S1: Statistical data of the tested models for each enzyme*. Compilation of all statistical data obtained for all tested models. Note – All above data was obtained through the anova() function and the r2glmm package. Underlined/bold models are the models that had the lowest AICc and were prioritized for data interpretations. DenDF – denominator degrees of freedom; R^2^M – marginal r-squared; R^2^C – conditional r-squared; CD – cestodes density; BD – black spots density; BC – body condition; * – interaction between factors and covariables; + – no interaction between factors and covariables.

## Literature Cited

Araújo, A., A.M. Jansen, F. Bouchet, K. Reinhard and L.F. Ferreira. 2003. Parasitism, the diversity of life, and paleoparasitology. Memórias do Instituto Oswaldo Cruz 98: 5–11.

Berra, T.M., and R.-J. Au. 1978. Incidence of black spot disease in fishes in Cedar Fork Creek, Ohio. Ohio J. Sci. 78(6):318.

Binning, S.A., D.G. Roche and C. Layton. 2013. Ectoparasites increase swimming costs in a coral reef fish. Biology letters 9: 20120927.

Binning, S.A., A.K. Shaw and D.G. Roche. 2017. Parasites and host performance: incorporating infection into our understanding of animal movement. Integrative and Comparative Biology 57: 267–280.

Bonneaud, C., J. Mazuc, G. Gonzalez, C. Haussy, O. Chastel, B. Faivre and G. Sorci. 2003. Assessing the cost of mounting an immune response. The American Naturalist 161: 367–379.

Bronte, V., and M.J. Pittet. 2013. The spleen in local and systemic regulation of immunity. Immunity 39: 806–818.

Caffaro, C.E., and J.C. Boothroyd. 2011. Evidence for host cells as the major contributor of lipids in the intravacuolar network of Toxoplasma-infected cells. Eukaryotic Cell 10: 1095–1099.

Capaldi, R.A. 1990. Structure and function of cytochrome c oxidase. Annual review of biochemistry 59: 569–596.

Careau, V., and T.G. Jr. 2012. Performance, personality, and energetics: correlation, causation, and mechanism. Physiological and Biochemical Zoology 85: 543–571.

Chabot, D., D. McKenzie and J. Craig. 2016. Metabolic rate in fishes: definitions, methods and significance for conservation physiology. pp. 1–9. Wiley Online Library.

Charron, A.J., and L.D. Sibley. 2002. Host cells: mobilizable lipid resources for the intracellular parasite Toxoplasma gondii. Journal of cell science 115: 3049–3059.

Cren, E.D.L. 1951. The length-weight relationship and seasonal cycle in gonad weight and condition in the perch (Perca fluviatilis). The Journal of Animal Ecology 20: 201.

Debeffe, L., P.D. Mcloughlin, S.A. Medill, K. Stewart, D. Andres, T. Shury, B. Wagner, E. Jenkins, J.S. Gilleard and J. Poissant. 2016. Negative covariance between parasite load and body condition in a population of feral horses. Parasitology 143: 983–997.

Dobson, A., K.D. Lafferty, A.M. Kuris, R.F. Hechinger and W. Jetz. 2008. Homage to Linnaeus: how many parasites? How many hosts? Proceedings of the National Academy of Sciences 105: 11482–11489.

Driedzic, W.R., J.R. Bailey and D.H. Sephton. 1996. Cardiac adaptations to low temperature in non □polar teleost fish. Journal of Experimental Zoology Part A: Ecological Genetics and Physiology 275: 186–195.

Escoll, P., L. Platon and C. Buchrieser. 2019. Roles of mitochondrial respiratory complexes during infection. Immunometabolism 1.

Fänge, R., and S. Nilsson. 1985. The fish spleen: structure and function. Experientia 41: 152–158.

Fischer, H., and R.S. Freeman. 1969. Penetration of parenteral plerocercoids of Proteocephalus ambloplitis (Leidy) into the gut of smallmouth bass. The Journal of Parasitology: 766–774.

Freitak, D., I. Ots, A. Vanatoa and P. Hörak. 2003. Immune response is energetically costly in white cabbage butterfly pupae. Proceedings of the Royal Society of London. Series B: Biological Sciences 270: S220–S222.

Gamperl, A.K., and A. Farrell. 2004. Cardiac plasticity in fishes: environmental influences and intraspecific differences. Journal of Experimental Biology 207: 2539–2550.

Guitard, J.J., E. Chrétien, J. De Bonville, D.G. Roche, D. Boisclair and S.A. Binning. 2022. Increased parasite load is associated with reduced metabolic rates and escape responsiveness in pumpkinseed sunfish. Journal of Experimental Biology 225.

Hiltunen, J.K., K.J. Autio, M.S. Schonauer, V.A.S. Kursu, C.L. Dieckmann and A.J. Kastaniotis. 2010. Mitochondrial fatty acid synthesis and respiration. Biochimica et Biophysica Acta (BBA) - Bioenergetics 1797: 1195–1202.

Holmstrup, M., A.-M. Bindesbøl, G.J. Oostingh, A. Duschl, V. Scheil, H.-R. Köhler, S. Loureiro, A.M. Soares, A.L. Ferreira and C. Kienle. 2010. Interactions between effects of environmental chemicals and natural stressors: a review. Science of the Total Environment 408: 3746–3762.

Hortová □Kohoutková, M., P. Lázničková and J. Frič. 2021. How immune □ cell fate and function are determined by metabolic pathway choice. BioEssays 43: 2000067.

Hugghins, E.J. 1959. Parasites of fishes in South Dakota. Bulletins 484.

Hunter-Manseau, F., V. Desrosiers, N.R. Le Francois, F. Dufresne, H.W. Detrich, 3rd, C. Nozais and P.U. Blier. 2019. From Africa to Antarctica: exploring the metabolism of fish heart mitochondria across a wide thermal range. Front Physiol 10: 1220.

Jakob, E.M., S.D. Marshall and G.W. Uetz. 1996. Estimating fitness: a comparison of body condition indices. Oikos: 61–67.

Kastaniotis, A.J., K.J. Autio, J.M. Keratar, G. Monteuuis, A.M. Makela, R.R. Nair, L.P. Pietikainen, A. Shvetsova, Z. Chen and J.K. Hiltunen. 2017. Mitochondrial fatty acid synthesis, fatty acids and mitochondrial physiology. Biochim Biophys Acta Mol Cell Biol Lipids 1862: 39–48.

Lagrue, C., and R. Poulin. 2015. Measuring fish body condition with or without parasites: does it matter? J Fish Biol 87: 836–847.

Laiz-Carrión, R., S. Sangiao-Alvarellos, J.M. Guzmán, M.P. Martín del Río, J.M. Míguez, J.L. Soengas and J.M. Mancera. 2002. Energy metabolism in fish tissues related to osmoregulation and cortisol action. Fish Physiology and Biochemistry 27: 179–188.

Laurent, P., S. Holmgren and S. Nilsson. 1983. Nervous and humoral control of the fish heart: structure and function. Comparative Biochemistry and Physiology Part A: Physiology 76: 525–542.

Lemieux, H., and P.U. Blier. 2022. Exploring thermal sensitivities and adaptations of oxidative phosphorylation pathways. Metabolites 12: 360.

Lemieux, H., P.U. Blier and E. Gnaiger. 2017. Remodeling pathway control of mitochondrial respiratory capacity by temperature in mouse heart: electron flow through the Q-junction in permeabilized fibers. Scientific reports 7: 1–13.

Lemly, A.D., and G.W. Esch. 1984. Effects of the trematode Uvulifer ambloplitis on juvenile bluegill sunfish, Lepomis macrochirus: ecological implications. The Journal of Parasitology 70: 475–492.

Leung, T.L., and R. Poulin. 2008. Parasitism, commensalism, and mutualism: exploring the many shades of symbioses. Vie et Milieu/Life & Environment: 107–115.

Lewin, R. 1982. Symbiosis and parasitism—definitions and evaluations. BioScience 32: 254–260.

Magistretti, P., and I. Allaman. 2013. Brain energy metabolism. pp. 1591–1620. Neuroscience in the 21st century: from basic to clinical. Springer New York.

Marcogliese, D.J. 2004. Parasites: small players with crucial roles in the ecological theater. EcoHealth 1: 151–164.

Margolis, L., and J.R. Arthur. 1979. Synopsis of the parasites of fishes of Canada. Bulletin of the Fisheries Research Board of Canada.

McElroy, E.J., and I. De Buron. 2014. Host performance as a target of manipulation by parasites: a meta-analysis. The Journal of Parasitology 100: 399–410.

Mills, E.L., B. Kelly and L.A.J. O’Neill. 2017. Mitochondria are the powerhouses of immunity. Nat Immunol 18: 488–498.

Nadler, L.E., E. Bengston, E.J. Eliason, C. Hassibi, S.H. Helland □ Riise, I.B. Johansen, G.T. Kwan, M. Tresguerres, A.V. Turner and K.L. Weinersmith. 2021. Brain □ A infecting parasite impacts host metabolism both during exposure and after infection is established. Functional Ecology 35: 105–116.

Nath, S., and J. Villadsen. 2015. Oxidative phosphorylation revisited. Biotechnology and Bioengineering 112: 429–437.

O’Dwyer, K., F. Dargent, M.R. Forbes and J. Koprivnikar. 2020. Parasite infection leads to widespread glucocorticoid hormone increases in vertebrate hosts: A meta □ analysis. Journal of animal ecology 89: 519–529.

Ots, I., A.B. Kerimov, E.V. Ivankina, T.A. Ilyina and P. Hõrak. 2001. Immune challenge affects basal metabolic activity in wintering great tits. Proceedings of the Royal Society of London. Series B: Biological Sciences 268: 1175–1181.

Pernas, L., C. Bean, J.C. Boothroyd and L. Scorrano. 2018. Mitochondria restrict growth of the intracellular parasite Toxoplasma gondii by limiting its uptake of fatty acids. Cell Metab 27: 886–897 e884.

Poulin, R. 2010. Parasite manipulation of host behavior: an update and frequently asked questions. pp. 151–186. Advances in the Study of Behavior. Elsevier.

R Development Core Team. 2022. R: A language and environment for statistical computing. R Foundation for Statistical Computing, Vienna, Austria.

Ryan, D.G., C. Frezza and L.A. O’Neill. 2021. TCA cycle signalling and the evolution of eukaryotes. Curr Opin Biotechnol 68: 72–88.

Salin, K., S.K. Auer, B. Rey, C. Selman and N.B. Metcalfe. 2015. Variation in the link between oxygen consumption and ATP production, and its relevance for animal performance. Proc Biol Sci 282: 20151028.

Sánchez, C.A., D.J. Becker, C.S. Teitelbaum, P. Barriga, L.M. Brown, A.A. Majewska, R.J. Hall and S. Altizer. 2018. On the relationship between body condition and parasite infection in wildlife: a review and meta □ analysis. Ecology letters 21: 1869–1884.

Santoro, M., S. Mattiucci, T. Work, R. Cimmaruta, V. Nardi, P. Cipriani, B. Bellisario and G. Nascetti. 2013. Parasitic infection by larval helminths in Antarctic fishes: pathological changes and impact on the host body condition index. Diseases of aquatic organisms 105: 139–148.

Schulz, H. 1991. Beta oxidation of fatty acids. Biochim Biophys Acta 1081: 109–120.

Shaw, D., and A. Dobson. 1995. Patterns of macroparasite abundance and aggregation in wildlife populations: a quantitative review. Parasitology 111: S111–S133.

Simon, A., D. Thomas, J. Blondel, P. Perret and M.M. Lambrechts. 2004. Physiological ecology of Mediterranean blue tits (Parus caeruleus L.): effects of ectoparasites (Protocalliphora spp.) and food abundance on metabolic capacity of nestlings. Physiological and Biochemical Zoology 77: 492–501.

Slavík, O., P. Horký, K. Douda, J. Velíšek, J. Kolářová and P. Lepič. 2017. Parasite-induced increases in the energy costs of movement of host freshwater fish. Physiology & behavior 171: 127–134.

Smith, P.e., R.I. Krohn, G.T. Hermanson, A.K. Mallia, F.H. Gartner, M. Provenzano, E.K. Fujimoto, N.M. Goeke, B.J. Olson and D. Klenk. 1985. Measurement of protein using bicinchoninic acid. Analytical biochemistry 150: 76–85.

Soengas, J.L., and M. Aldegunde. 2002. Energy metabolism of fish brain. Comparative Biochemistry and Physiology Part B: Biochemistry and Molecular Biology 131: 271–296.

Stanley, I.A., S.M. Ribeiro, A. Giménez-Cassina, E. Norberg and N.N. Danial. 2014. Changing appetites: the adaptive advantages of fuel choice. Trends in cell biology 24: 118–127.

Stavru, F., F. Bouillaud, A. Sartori, D. Ricquier and P. Cossart. 2011. Listeria monocytogenes transiently alters mitochondrial dynamics during infection. Proceedings of the National Academy of Sciences 108: 3612–3617.

Suzuki, M., O. Danilchanka and J.J. Mekalanos. 2014a. Vibrio cholerae T3SS effector VopE modulates mitochondrial dynamics and innate immune signaling by targeting Miro GTPases. Cell host & microbe 16: 581–591.

Suzuki, N., R.M. Rivero, V. Shulaev, E. Blumwald and R. Mittler. 2014b. Abiotic and biotic stress combinations. New Phytologist 203: 32–43.

Thibault, M., P. Blier and H. Guderley. 1997. Seasonal variation of muscle metabolic organization in rainbow trout (Oncorhynchus mykiss). Fish Physiology and Biochemistry 16: 139–155.

Timi, J.T., and R. Poulin. 2020. Why ignoring parasites in fish ecology is a mistake. Int J Parasitol 50: 755–761.

Turner, W.C., W.D. Versfeld, J.W. Kilian and W.M. Getz. 2012. Synergistic effects of seasonal rainfall, parasites and demography on fluctuations in springbok body condition. Journal of animal ecology 81: 58–69.

Wuenschel, M.J., W.D. McElroy, K. Oliveira and R.S. McBride. 2019. Measuring fish condition: an evaluation of new and old metrics for three species with contrasting life histories. Canadian Journal of Fisheries and Aquatic Sciences 76: 886–903.

