## Supplemental Table 1 for "Mitochondrial metabolism and body condition of naturally infected sunfish (*Lepomis gibbosus*)"

|  | Model | Factor/covariable | DenDF | F-value | P-value | R^2^M | R^2^C |
| --- | --- | --- | --- | --- | --- | --- | --- |
| CCO | CD*organ | CD  Organ  CD*organ | 20  60  60 | 0.88  324.45  2.56 | 0.36  < 2e-16  0.064 | 0.95 | 0.96 |
|  | BD*organ | BD  Organ  BD*organ | 20  60  60 | 1.83  240.19  0.99 | 0.19  < 2e-16  0.40 | 0.94 | 0.95 |
|  | **BC*organ** | BC  Organ  BC*organ | 20  60  60 | 4.08  8.04  3.00 | 0,057  0,00018  0,038 | 0,95 | 0,96 |
| CS | CD*organ | CD  Organ  CD*organ | 2  60  60 | 0.49  254.91  0.42 | 0.49  < 2e-16  0.74 | 0.87 | 0.95 |
|  | BD*organ | BD  Organ  BD*organ | 20  60  60 | 0.19  203.96  0.29 | 0.66  < 2e-16  0.83 | 0.87 | 0.95 |
|  | BC*organ | BC  Organ  BC*organ | 20  60  60 | 1.44  7.29  0.95 | 0.24  0.00030  0.42 | 0.87 | 0.95 |
|  | **BC+organ** | BC  Organ | 20  63 | 1.44  531.45 | 0.24  < 2e-16 | 0.87 | 0.95 |
| CPT | CD*organ | CD  Organ  CD*organ | 20  60  60 | 0.063  36.08  0.99 | 0.80  1.868e-13  0.40 | 0.62 | 0.71 |
|  | BD*organ | BD  Organ  BD*organ | 20  60  60 | 0.091  17.97  0.54 | 0.77  1.946e-08  0.66 | 0.61 | 0.70 |
|  | BC*organ | BC  Organ  BC*organ | 20  60  60 | 3.91  2.61  1.19 | 0.062  0.059  0.32 | 0.64 | 0.71 |
|  | **BC+organ** | BC  Organ | 20  63 | 3.91  60.91 | 0.062  < 2e-16 | 0.64 | 0.71 |
| ETS | CD*organ | CD  Organ  CD*organ | 20  60  60 | 3.12  2.51  0.082 | 0.093  0.068  0.97 | 0.18 | 0.37 |
|  | BD*organ | BD  Organ  BD*organ | 20  60  60 | 1.13  2.32  0.14 | 0.30  0.085  0.93 | 0.15 | 0.37 |
|  | BC*organ | BC  Organ  BC*organ | 20  60  60 | 0.096  0.91  1.02 | 0.76  0.44  0.39 | 0.15 | 0.40 |
|  | **CD+organ** | CD  Organ | 2  63 | 3.12  6.36 | 0.093  0.00078 | 0.16 | 0.38 |

Note – All above data was obtained through the anova() function and the r2glmm package. Underlined/bold models are the models that had the lowest AICc and were prioritized for data interpretations. DenDF – denominator degrees of freedom; R^2^M – marginal r-squared; R^2^C – conditional r-squared; CD – cestodes density; BD – black spots density; BC – body condition
