## Supplementary figures and images for "Mitochondrial metabolism and body condition of naturally infected sunfish (*Lepomis gibbosus*)"

### Supplemental Figure 1

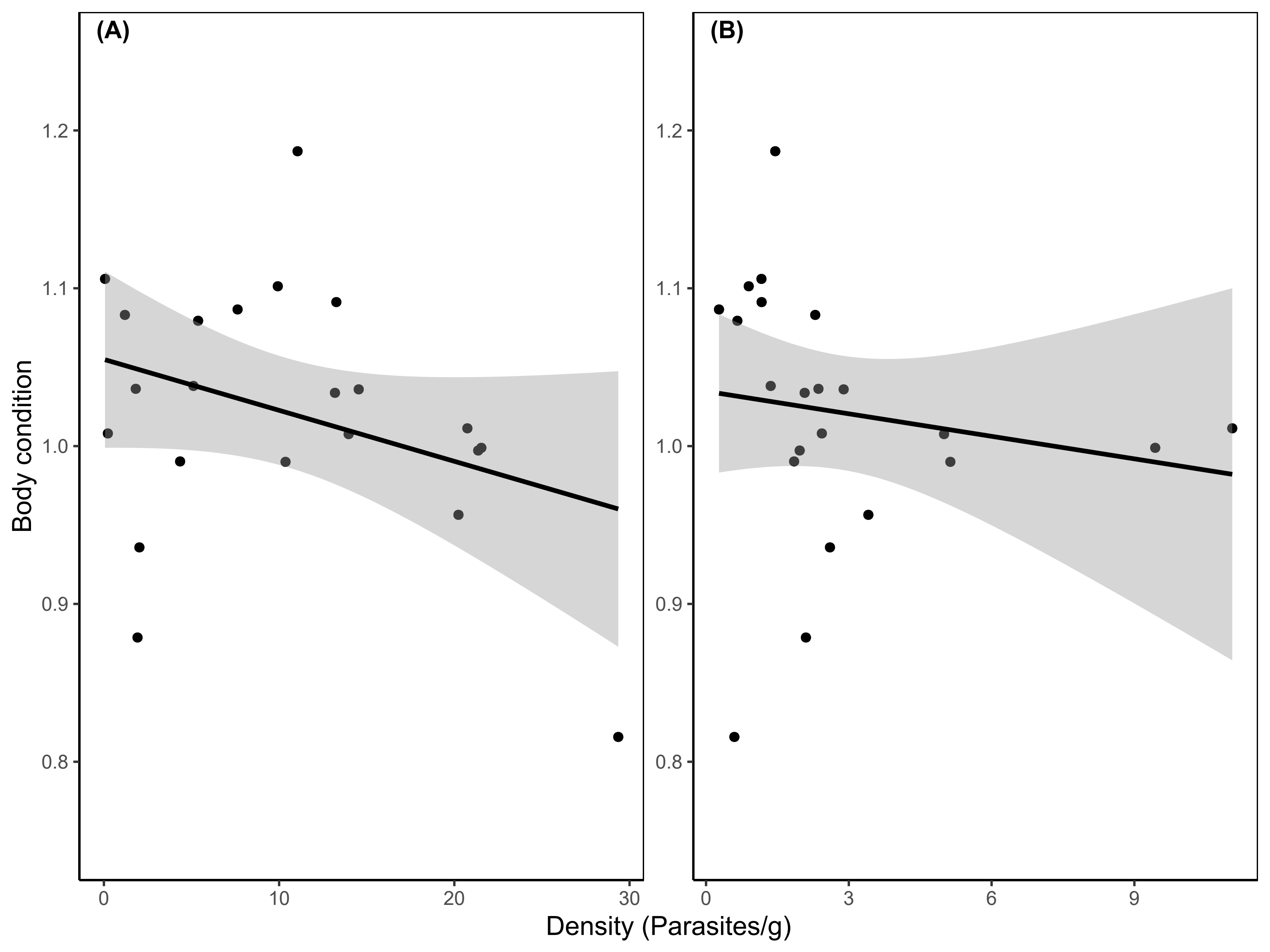
